# Energy content of anchovy and sardine using surrogate calorimetry methods

**DOI:** 10.1101/2021.02.01.429042

**Authors:** Claudia Campanini, Marta Albo-Puigserver, Sara Gérez, Elena Lloret-Lloret, Joan Giménez, Maria Grazia Pennino, Jose Maria Bellido, Ana I. Colmenero, Marta Coll

## Abstract

European anchovy (*Engraulis encrasicolus*) and sardine (*Sardina pilchardus*) are crucial species for the marine ecosystem of the Northwestern Mediterranean Sea. They account for a high percentage of fish landings and they represent an important economic income. Concerns over their stock status are rising in recent years as biomass, growth, reproductive capacity and body condition of both species are declining. Therefore, there is an urgent need for a continuous and fast body condition monitoring scheme. Energy storage variability has important implications for both fish recruitment and population structure. Direct condition indices, such as bomb calorimetry, are highly reliable for measuring the energy content, but extremely time-consuming. Alternatively, fatmeter analysis and relative condition index (Kn) have been proposed as effective indirect methods. The aim of this study is to test the application of fatmeter as a surrogate of bomb calorimetry to infer the energy content of both small pelagic fishes. For the validation, fatmeter values were compared with both energy density (ED; via bomb calorimetry) and Kn values. Individuals of both species were sampled monthly in Barcelona harbor for a year in order to assess seasonal variations in energy content. Our results highlight that fatmeter measurements are strongly correlated with calorimetry for sardine, while a weaker but significant correlation was found for anchovy. The observed differences between the two species are related to their breeding strategies. Based on this study, fatmeter analysis appears to be a faster and suitable method to evaluate the energy content of both species routinely., In addition, we provide a linear model to infer ED from fatmeter values of both small pelagic fish. Eventually, these findings could allow for the avoidance of bomb calorimetry and could be used to implement body condition monitoring protocols, and to boost continuous large-scale monitoring.

## 1. Introduction

Fish condition has been defined as a good indicator of health and physiological status of fish (Lloret et al., 2012, 2014). Although there is not a univocal definition of fish condition, it is frequently considered as a measure of individual energy storage (Gatti et al., 2018a; McPherson et al., 2011). Energy content of fish is a dynamic measure of their energy balance, which is regulated by physiological functions (i.e. growth, maintenance or reproduction) and environmental factors (i.e. temperature and food availability) (Albo-Puigserver et al., 2017). As a result, energy content of fish species has also been used to track changes in the ecosystem and as proxy of environmental status, especially of species that play a pivotal role for the functioning of the ecosystem such as small pelagic fish (Nikolsky et al., 2012; Shulman et al., 2005).

Since energy content influences growth rate, reproduction success and mortality rate, the assessment of body condition combined with other variables (e.g. biomass, exploitation, abundance) is useful to monitor the state of fish stocks and to predict future changes in terms of abundance and productivity (Lambert and Dutil, 1997; Lloret et al., 2012). Moreover, energy condition indices such as energy density (ED) are needed to build bioenergetic models (Gatti et al., 2017), which allow the quantification of energy allocation and can be applied in physiology, ecology, aquaculture, and fisheries management (Deslauriers et al., 2017).

In order to evaluate fish condition, direct and indirect methods are currently used (Schloesser and Fabrizio, 2017; Lloret et al., 2014). Direct indices, such as ED from direct calorimetry or lipid content, provide a precise measure of energy storage that can be used as an estimation of fitness (Lloret et al., 2014), but are highly time-consuming and not suitable for a continuous monitoring. By contrast, indirect indices, which can be based on length-mass relationship (length-based indices), organ mass (i.e. organosomatic indices), or tissue properties (e.g. fatmeter) (Schloesser and Fabrizio, 2017), allow a rapid assessment of fish condition. However, prior to their use as an indirect proxy of energy storage, they should be validated investigating their relationship with direct condition indexes (Davidson and Marshall, 2010; McPherson et al., 2011; Schulte-Hostedde et al., 2005).

When evaluating the variability in body condition, other parameters, such as size, life stage (reproductive/resting period) and season need to be taken into consideration (Brosset et al 2015, Gatti et al., 2018). Indeed, energy content is strongly affected by temporal changes in food availability and this in turns impacts life-history strategy (Albo-Puigserver et al., submitted; Lloret et al., 2014). This is the case of European anchovy *Engraulis encrasicolus* (Linnaeus, 1758) and sardine (also named European pilchard) *Sardina pilchardus* (Walbaum, 1792), which display different energy allocation and life-history strategies. European anchovy (hereinafter anchovy) is an income breeder that allocates energy directly to reproduction, which occurs in spring and summer; conversely, sardine is a capital breeder that stores energy during the resting period prior to the reproduction in autumn and winter (Albo-Puigserver et al., 2017; Costalago and Palomera, 2014; Ganias et al., 2007; McBride et al., 2015; Palomera et al., 2007).

Sardine and anchovy stocks in the western Mediterranean Sea are currently highly fished or overexploited (Coll and Bellido, 2019; FAO, 2018; GFCM, 2019). In the last decades, a decrease in body condition has been observed for both species in several parts of the Mediterranean Sea, with a concurrent drop in landings, abundance, growth and age structure (Albo-Puigserver et al. submitted, Brosset et al., 2017; Coll and Bellido, 2019; Van Beveren et al., 2014). Due to their importance in biomass in mid-trophic positions, sardine and anchovy are key elements of the marine food web having an essential role in the energy transfer from lower to upper trophic levels (Coll et al., 2008, 2006; Cury et al., 2000, 2011; Piroddi et al., 2015). Thus, a decline in fish condition can have a bottom-up effect on top-predators populations negatively impacting their breeding success and fitness (Crawford et al., 2006; Österblom et al., 2008; Piroddi et al., 2017).

In order to evaluate the body condition of sardine and anchovy, morphometric indices, such as Le Cren relative condition index (Kn), have been widely used (Brosset et al., 2015b; Van Beveren et al., 2014). This index rely on the assumption that any deviation from the weight-length relationship standard can be assumed as an indication of the relative fitness of the individual (Lloret et al., 2014). Being low-cost, quick, and usually non-disruptive, these morphometric indices are used in many fields. Although their use have been validated for sardine and anchovy (Albo-Puigserver et al., 2020; Brosset et al., 2015a; Gatti et al., 2018a; McPherson et al., 2011), their utilization to infer energy content is still controversial (Davidson and Marshall, 2010; McPherson et al., 2011; Sardenne et al., 2016; Stevenson and Woods Jr, 2006).

On the other hand, direct measures of energy density using bomb calorimetry, defined as the energy content per weight unit (kJ·g^−1^), have demonstrated to be a precise measure of fish bioenergetics condition (Albo-Puigserver et al., 2020, 2017; Gatti et al., 2018a; Tirelli et al., 2006). The benefit of bomb calorimetry is that it gives information on the average energy of the proximate composition of fish (weighted average of protein, lipid and carbohydrates energy densities) (Gatti et al., 2018a). Unfortunately, a continuous monitoring of body condition with this technique is impractical as it is time-consuming, costly, and impossible to carry in-situ (i.e. monitoring in harbors or in oceanographic campaigns).

The use of fish fatmeter has many advantages as the measurement is rapid, repeatable, accurate, non-destructive, and the device is portable, making it very convenient for field research. Fish store energy mainly as lipids and thus they drive most of the variation of energy density measurements (Anthony et al., 2000; Rosa et al., 2010; Spitz et al., 2010). Numerous studies described a positive correlation between fatmeter and lipid content or morphometric condition indices (Bayse et al., 2018; Brosset et al., 2015a; Davidson and Marshall, 2010; Másílko et al., 2016). Furthermore the accuracy of fatmeter as a surrogate of direct techniques to estimate energy reserves has been assessed for several species (Bayse et al., 2018; Goñi and Arrizabalaga, 2010; Mann et al., 2009; Mesa and Rose, 2015; Schloesser and Fabrizio, 2017). As for the sardine and anchovy, the use of Distell Fish Fatmeter has been validated as a good indirect method to estimate lipid content (Brosset et al., 2015a), but whether it could also provide accurate estimates of energy content has not been investigated. Therefore, the main objective of this study was to assess the suitability of fish fatmeter as surrogate of bomb calorimetry to estimate energy density and therefore fish condition of anchovy and sardine from the Northwestern Mediterranean Sea, considering temporal and ontogenetic changes in body condition. Furthermore, we compared fatmeter measurements with Le Cren relative condition index, as this latter index has already been validated for both species as a surrogate for energy content (Brosset et al., 2015a). Secondly, where the correlation between fatmeter values (expressed in lipid percentage) and energy density (ED; expressed in kJ·g^−1^) turned out to be strong, a model fitted to this relationship was provided, as such model could be used to infer ED from fatmeter values allowing a faster body condition assessment protocol for sardine and anchovy in the future. Moreover, the inferred ED values could be used to measure the transfer of energy between compartments and ultimately for bioenergetic (Gatti et al., 2017; Pecquerie et al., 2009) and ecosystem models.

## 2. Material and methods

### 2.1. Study area and sampling

The present study was carried off Barcelona harbor in the Catalan Sea, Northwestern Mediterranean Sea (Figure 1). Overall, the Catalan coast presents a relatively narrow continental shelf (Martín et al., 2008). The hydrography of this coastal area is highly influenced by anthropogenic factors due to the proximity of the city and its high urban density (Romero et al., 2014). Salinity increases moving towards the edge of the continental shelf due to the high freshwater discharge of both natural and anthropogenic origin in the area (Guillén et al., 2019). Indeed, freshwater inputs include not only discharge from the Besòs River to the north and the Llobregat River to the south, but also from many storm sewers (Figure 1). Such inputs consist mainly of treated waters, however even untreated waters are released to the sea during heavy rainfall (Arin et al., 2013). In temperate oceanic zones, such as this area, primary production displays significant interannual changes (Bosc et al., 2004; Estrada, 1996). The observed peaks in phytoplankton biomass (main peak in winter-spring and secondary peak in autumn) can be explained by both increased freshwater discharges, related to higher precipitations, and vertical mixing due to decreased temperatures and strong winds. Conversely, primary production reaches its minimum during summer due to the scarcity of nutrients caused by water stratification (Arin et al., 2013).

**Figure 1.**
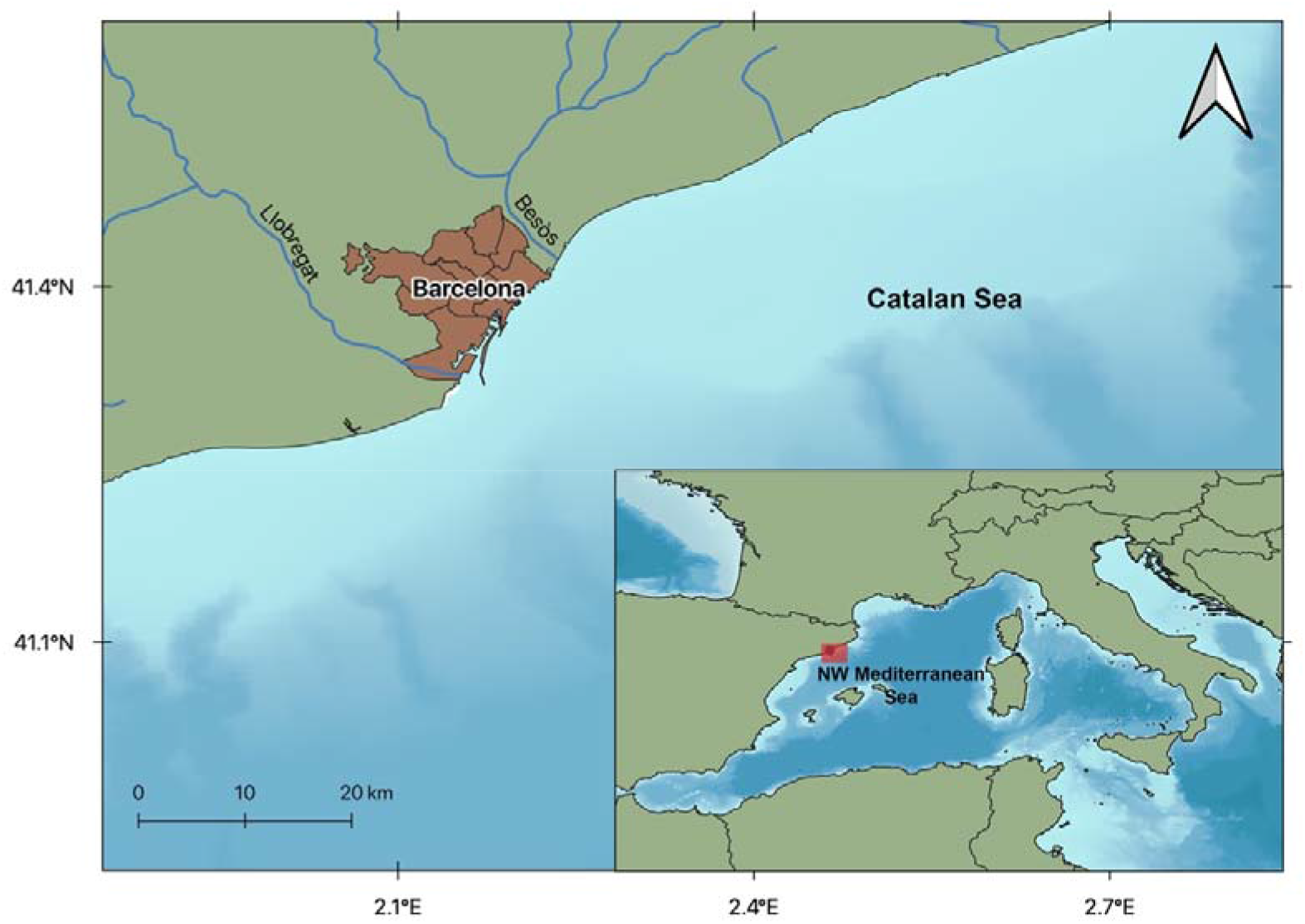
Study area corresponding to the coast of Barcelona where samples were collected.

Samples for both species were obtained monthly, from July 2018 to June 2019, from commercial purse-seiners landings off the Barcelona harbor. Data of anchovy from November and December 2019 are missing due to lack of catch (November) and anchovy annual fishing closure (December). For all the statistical analyses, seasons were defined as follows: summer from July to September, autumn from October to December, winter from January to March, and spring from April to June. In order to capture length variability, 40 individuals covering all available targeted size-ranges were sampled for each species per month. Size (cm), weight (g), sex (female, male and indeterminate), and fatmeter measurements (explained later in more detail) were recorded for each individual. Ten individuals belonging to different size ranges from each month were randomly selected for calorimeter analysis in order to obtain ED values from all the size-spectrum.

**Table 1.** Sample size by month and species for each index: Kn, fatmeter (Fatm) and energy density (ED). Minimum (min) and maximum (max) monthly values of body length (cm) during 2018-2019.

| Year | Month | Anchovy |  |  |  | Sardine |  |  |  |
| --- | --- | --- | --- | --- | --- | --- | --- | --- | --- |
|  |  | Body length<br>(cm)<br>(min-max) | Kn | Fatm | ED | Body length<br>(cm)<br>(min-max) | Kn | Fatm | ED |
| 2018 | July | 11.40-16.00 | 40 | 40 | 10 | 14.20-18.50 | 38 | 40 | 10 |
|  | August | 9.00-15.20 | 40 | 40 | 10 | 9.40-18.30 | 40 | 40 | 10 |
|  | September | 9.10-15.10 | 40 | 40 | 10 | 13.70-18.20 | 40 | 40 | 10 |
|  | October | 8.30-16.10 | 40 | 40 | 10 | 13.20-18.60 | 40 | 40 | 10 |
|  | November | - | - | - | - | 10.20-16.40 | 40 | 40 | 10 |
|  | December | - | - | - | - | 11.90-17.60 | 40 | 40 | 10 |
| 2019 | January | 12.00-15.70 | 40 | 35 | 10 | 12.00-18.20 | 40 | 40 | 10 |
|  | February | 10.30-15.30 | 40 | 40 | 10 | 9.90-18.30 | 40 | 40 | 10 |
|  | March | 12.70-15.20 | 40 | 40 | 10 | 12.30-16.20 | 40 | 40 | 10 |
|  | April | 10.50-15.00 | 40 | 40 | 9 | 13.20-16.80 | 40 | 40 | 10 |
|  | May | 10.60-14.50 | 40 | 40 | 10 | 12.50-16.10 | 40 | 40 | 10 |
|  | June | 11.30-15.00 | 40 | 40 | 10 | 12.50-16.20 | 40 | 40 | 10 |

### 2.2. Fatmeter analyses

The Fish fatmeter works exploiting the transmission of low-power microwaves to measure the dielectric properties and thus the water content of the tissue beneath the skin (Kent, 1990). We used an FFM-992 Distell Fatmeter, the so-called “Small Head Model”, to process the fatmeter samples, as the 3 cm-long sensor is more suitable for small-sized fish. Prior to the utilization, the instrument was calibrated using the standard calibration provided by the manufacturer, i.e. ANCHOVY-2 and SARDINE-2. Two measures were taken for each side of the fish placing the microstrip sensor along the lateral line, as recommended in the user manual (Distell, 2010). Finally, mean fatmeter values were calculated from the 4 measurements recorded for each specimen (2 for each side) (Brosset et al., 2015a). Some anchovy shorter than 10.3 cm were too small and the Fish Fatmeter could not record any measurement, so two individuals placed side by side were jointly measured and the same fatmeter value was recorded.

### 2.3. Energy density analyses

A total of 120 and 100 calorimetric analyses were performed for sardine and anchovy, respectively. Samples for direct calorimetry analysis were prepared following the protocol described by Albo-Puigserver et al. (2017) except for the oven temperature that was 60 °C.

ED was estimated as the heat release from the combustion of a small subsample and it was measured using Parr 6725 Semimicro Oxygen Bomb Calorimeter. The instrument was previously calibrated combusting benzoic acid according to the Operating Instruction Manual (Parr Instrument Company, 2012). Two 50-150 mg pellets per specimen were combusted in the calorimeter. In the eventuality that the two measurements differed by more than 5%, a third pellet from the same individual was analyzed. Values, expressed in calories per gram (cal·g^−1^) of dry weight (DW), were converted to kilojoules per gram of wet weight (kJ·g^−1^ WW). The conversion to a wet weight (WW) basis was achieved by multiplying mean ED values (kJ·g^−1^ DW) by dry weight proportion (dry weight proportion = dry weight · wet weight^−1^). Finally, mean ED of each individual was calculated from the subsamples’ measurements (Albo-Puigserver et al., 2017).

### 2.4. Relative condition index (Kn)

The relative condition index (Kn) was determined for each individual as the ratio of its total weight (TW) to the predicted weight (PW) for a fish of the same total length (Le Cren, 1951).

The predicted weight (PW) was computed using the weight–length regression expressed by the following equation: PW=a·TL^b^ (TL = Total Length).

The weight-length relationship is species- and population-specific and thus it was computed *ad hoc* for this study. The two coefficients, *a* and *b*, where estimated for the two species taking into account all the individuals from the sampling period (anchovies: a = 0.0036, b = 3.216; sardines: a = 0.0031, b = 3.334).

### 2.5. Statistical analyses

We explored the relationship of fatmeter values and ED with size (fish total length, in cm), using Generalized Additive Models (GAM). GAMs were run with the function *gam* included in R library *mgcv* (Wood, 2011) using restricted maximum likelihood (REML) for smoothing parameter estimation, while the R library *visreg* (Breheny and Burchett, 2017) was used for GAMs visualization. A Gaussian distribution with an identity link was used to fit condition indices. Since condition indices are influenced by many biological and environmental variables, GAMs were fitted for each species considering also sex and season and their concurrent effects. In particular, season and sex were included in GAMs as factors, while size was treated as a continuous variable. Finally, as the rest of the variability could be due to some individual intrinsic aspects, we also added a random individual effect, hereafter defined as ID. In order to avoid overfitting, degrees of freedom were restricted to 6 for both main effects and interactions.

GAMs by species were selected as the best-to-fit model based on the explained deviance and the Akaike Information Criterion (AIC) (Burnham and Anderson, 2002).

We also generated linear regression and log-log regression models of fatmeter values against DW. Furthermore, the relationship between DW and ED in wet weight basis (kJ·g^−1^ ww) described by Hartman and Brandt (1995) was investigated for each species as to compare our results with previous literature, i.e. Gatti et al., (2018) for both species in the Bay of Biscay and the English Channel, Albo-Puigserver et al. (2020; unpub. data) for both species in Tarragona coast and Tirelli et al. (2006) for anchovy in the Adriatic Sea. In particular, linear regression and log-log regression models were performed for each species between DW and ED. Then, results of the mentioned published studies were plotted against our results for visual comparison and the estimated regression parameters were analytically compared with the Welch-test. In addition, the relationships between percentage of DW and ED in dry weight basis (kJ·g^−1^ dw) for both species were plotted and are included in the Supplementary materials (Figure S1).

In order to assess the correlation and therefore the eventual interchangeability between indices, pairwise Spearman’s rank non-parametric correlation test was performed between Kn, fatmeter values and ED. We expected a large year-round variation in condition indices related to the reproduction investment; therefore, we split the original dataset according to the estimated reproductive and resting period for both species, the former lasting from October to March for sardine and from April to September for anchovy.

Finally, as we hypothesized that fatmeter values could allow indirect estimation of energy density, linear regression and log-log regression models were also performed to assess the relationship between fatmeter and ED for both species. Prior to fit the models, the presence of outliers was checked through the Bonferroni Outlier Test and they were eventually excluded. For the present study, all the statistical analyses were performed with R version 3.6.1 (R Core Team, 2019) and a p-value threshold of 5% was considered as significance level.

## 3. Results

### 3.1. Monthly variation in indirect and direct condition indices

Overall, sardine exhibited a higher seasonality in condition indices compared to anchovy, but in both species the lowest values of Kn, fatmeter and ED were found from November to February 2018-2019 (Figure 2). As for sardine, the percentage of lipid estimated by fatmeter measurements changed markedly throughout the year (Figure 2A). The highest mean values were recorded in July (18.00±1.64 lipid %) and followed by a slow decrease from August to October. Lipid percentage drastically declined in November and reached its minimum average value in December. A slow recovery was registered in late winter while from March onwards it surpassed the 10%.

**Figure 2.**
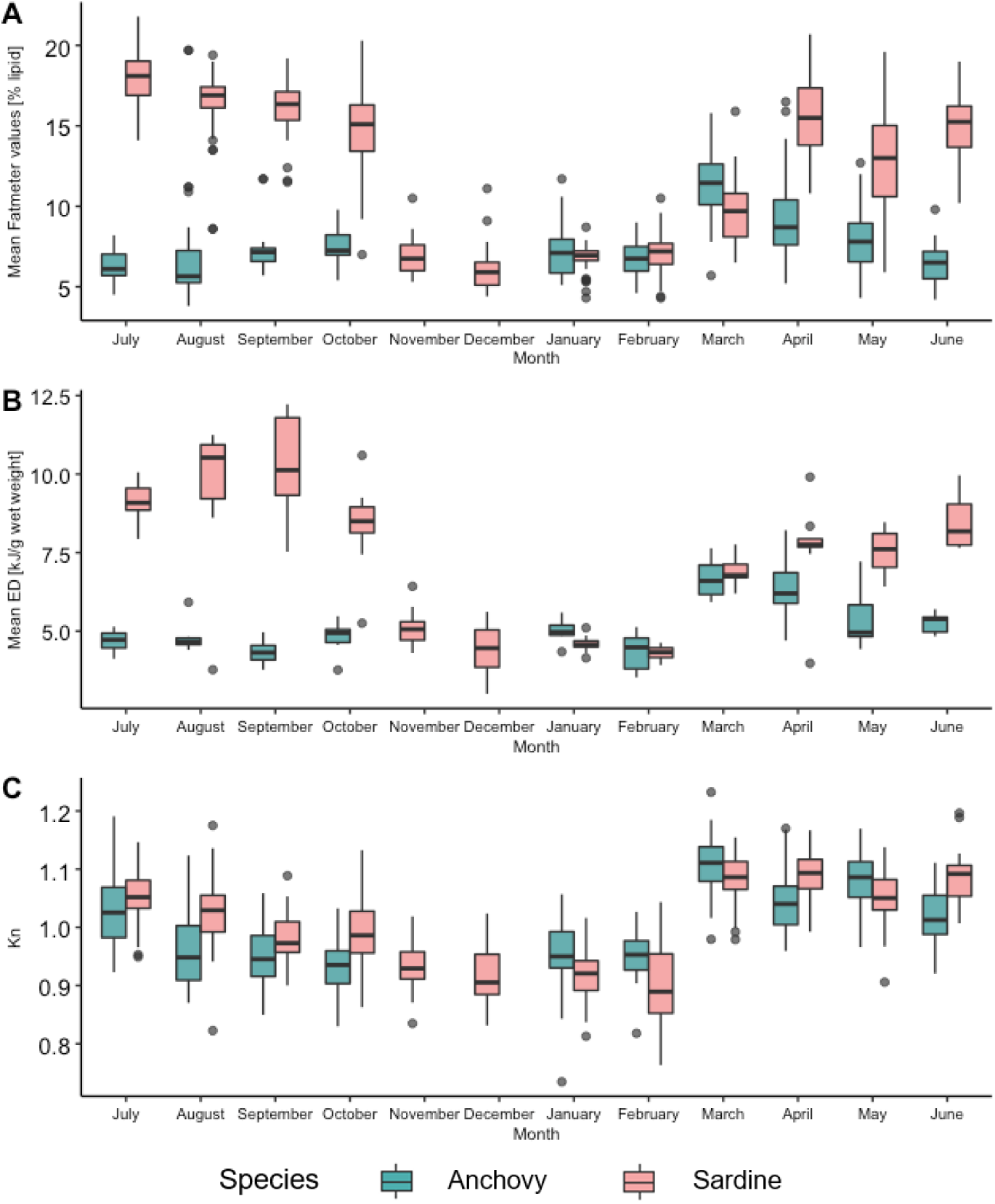
Boxplot of monthly variation of the three condition indices: A) Fatmeter, B) Energy density (ED), and C) Relative body condition (Kn), for anchovy (in dark cyan) and sardine (in coral rose) from July 2018 to June 2019. Boxplots hinges represent the 1^st^ and 3^rd^ quartiles. Horizontal lines represent the medians.

Regarding anchovy, the lipid percentage as measured with fatmeter was more stable, with mean values ranging from 5 to 10% all year round, apart from March when the mean value was higher (11.4±1.85 lipid %).

Months with relatively high ED seemed to have higher intra-monthly variability for both anchovy and sardine. Sardine ED was very high in late summer, reaching its peak in September, then started to fall from October and progressively rise again in March. Year-round ED pattern for anchovy was similar to its fatmeter pattern. Mean ED values oscillated around 5 kJ·g^−1^ until February with an increase between March and May, while they ranged from 5 to 7.5 kJ·g^−1^ during the rest of the year (Figure 2B).

Concerning Kn values for sardine, we observed a gradual decrease from July to February (February, 0.89±0.07) followed by a sharp rise in March (1.09±0.04). Regarding anchovy, lower monthly variation in Kn was observed and, similar to sardine, they displayed a substantial increase in March (Figure 2C).

### 3.2. Variation in energy content with length, sex, and season

For anchovy, the deviance explained by GAMs models fitted to the evolution of ED and fatmeter values with size (Figure 3) was 34.3% for fatmeter values and 52.7% for ED. Regarding fatmeter GAMs, all the effects proved to be significant (Table S2). However, the interaction between sex and season was not included in the model as its effect was only significant for females and males in winter and its inclusion did not substantially improve the explained deviance. Fatmeter GAMs by season were quite wiggly at visualization (Figure 3). Indeed, fatmeter values tended to increase with size in winter and spring but this trend was reversed in summer and autumn, as higher values were recorded for small-sized individuals. By contrast, the relationships found with models fitted to ED by season were linear and we excluded both sex and the interaction between sex and season from the model as they were not significant. ED GAM showed a rather slight increase in summer and autumn values in response to an increase in size, while they displayed a steeper linear slope in winter and spring. The lack of specimens shorter than 10 cm and longer than 15 cm during winter and spring explained the large confident interval in both ED and fatmeter models. Finally, the random individual effect was statistically relevant for both ED and fatmeter models.

**Figure 3.**
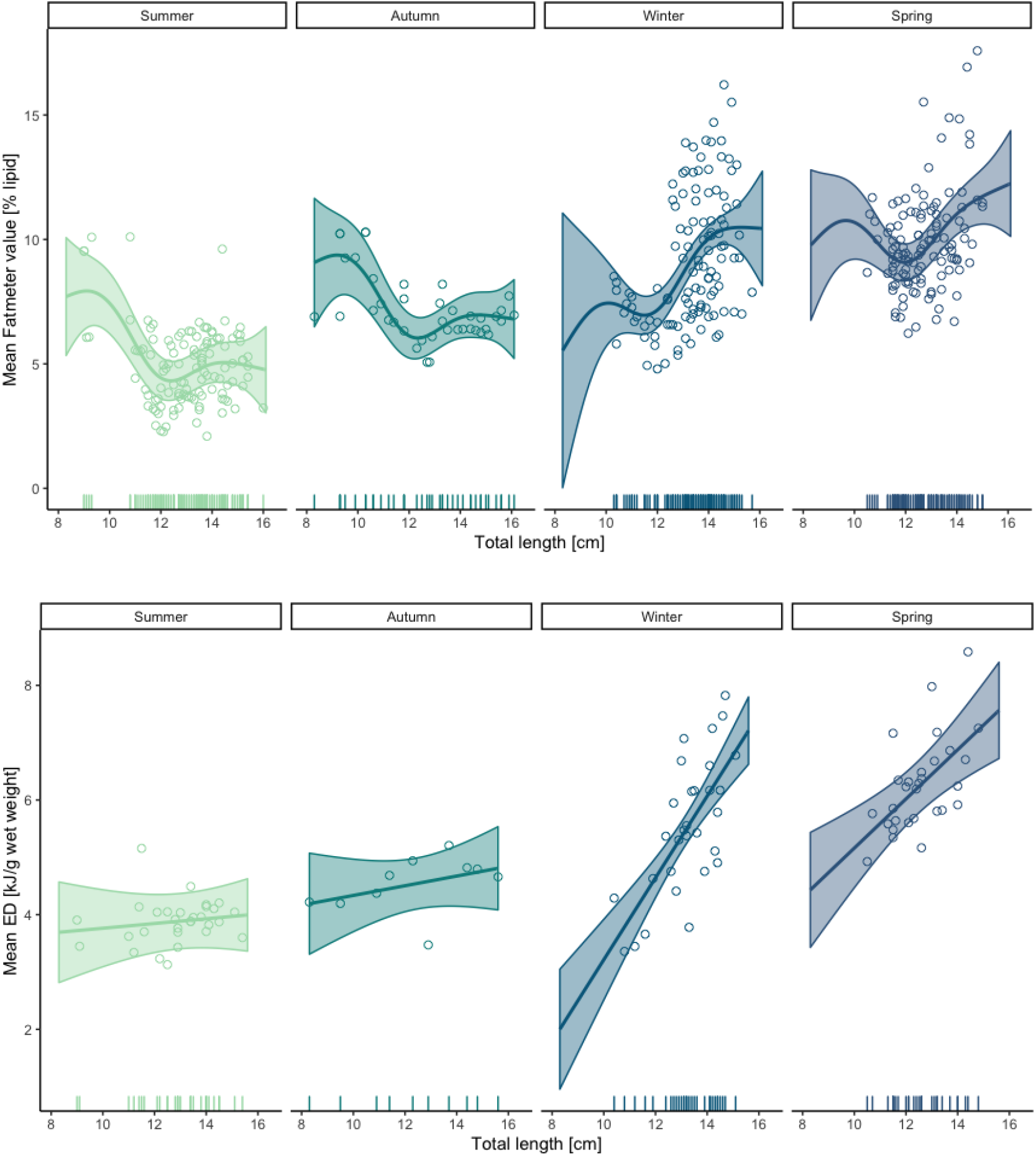
Partial effect plots of season on predicted fatmeter (top) and ED (bottom) values by the GAMs against total length for anchovy. Seasonal model fit with average predictions (continuous lines) and the 95% confidence intervals.

With regards to sardine, GAM fitted to the variability of fatmeter against total length, season, and sex explained 74.1% of the deviance, while the one fitted to ED explained the 69.2% (Table S3 and Figure 4). In accordance with the principle of parsimony, we excluded sex and its interaction with season from fatmeter model as their effects were not relevant, but for female-male difference in winter. Conversely, difference between sexes proved to be significant and therefore it was included in ED GAM. Notably, fatmeter values and size were not always positively related in all seasons; indeed, for summer and winter, increasing fatmeter values were estimated until a total length of 16 cm, while over this length the estimated fatmeter values tended to gradually decrease (Figure 4). ED GAMs showed that ED values increased with fish length during all the seasons with an almost linear trend. For both ED and fatmeter models in summer and spring, the large confidence intervals estimated for short individuals were consequent of the low number of specimen shorter than 13 cm collected during these seasons. The smoothed random individual effect exerted a significant influence only in fatmeter values variation.

**Figure 4.**
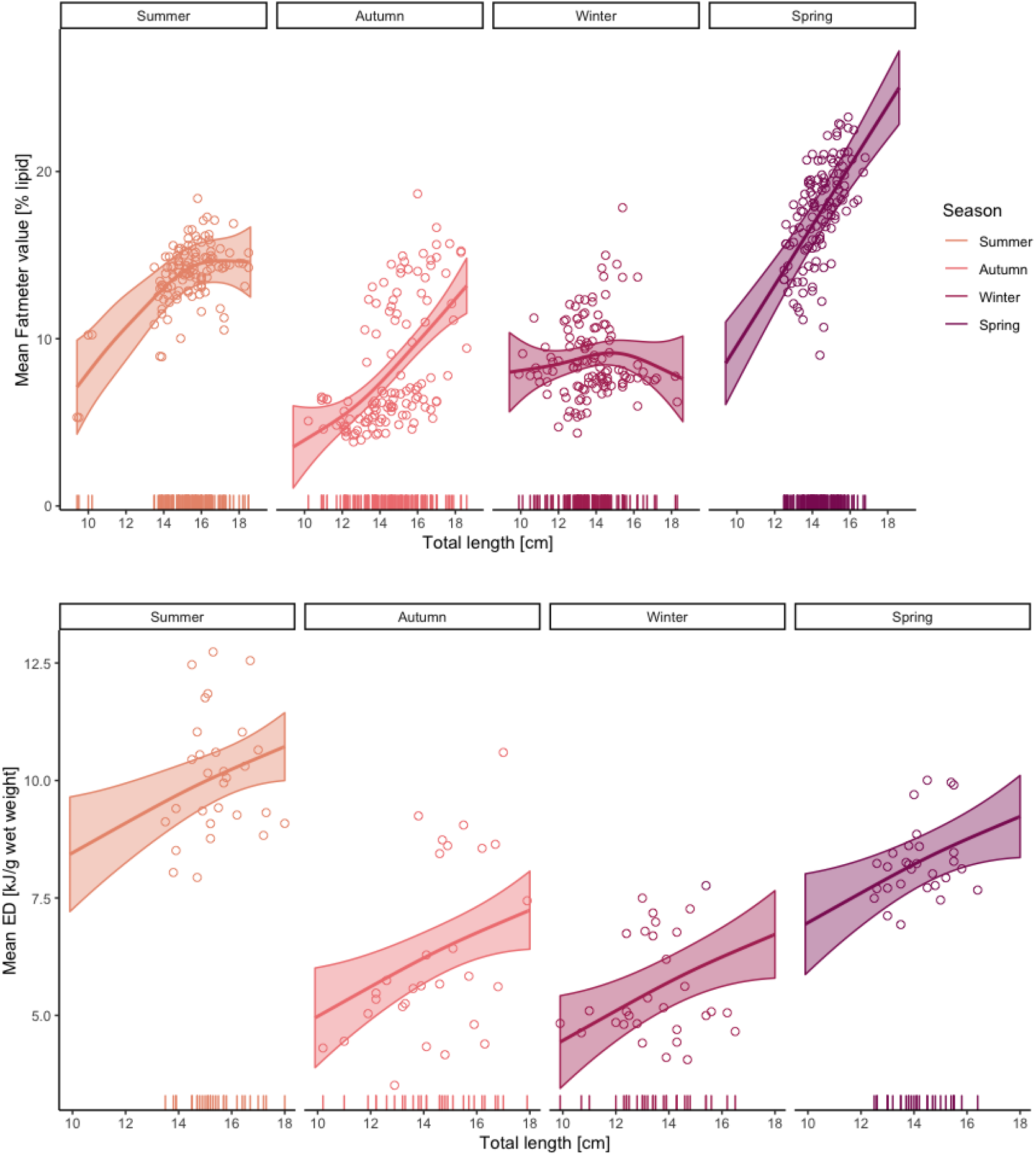
Partial effect plots of season on predicted fatmeter (top) and ED (bottom) values by the GAMs against total length for sardine. Seasonal model fit with average predictions (continuous lines) and the 95% confidence intervals.

Overall, GAMs fitted both to fatmeter and ED have rather high residual standard deviations for both anchovy (fatmeter GAM: RSD ≈ 1.9; ED GAM: RSD ≈ 0.75) and sardine (fatmeter GAM: RSD ≈ 2.5; ED GAM: RSD ≈ 1.30).

### 3.3. Fatmeter validation

Linear regression and log-log models showed that an increase in dry weight (DW) corresponded to an increase in both fatmeter values and energy density (ED) for both species, thus they were positively correlated (Figures 5).

**Figure 5.**
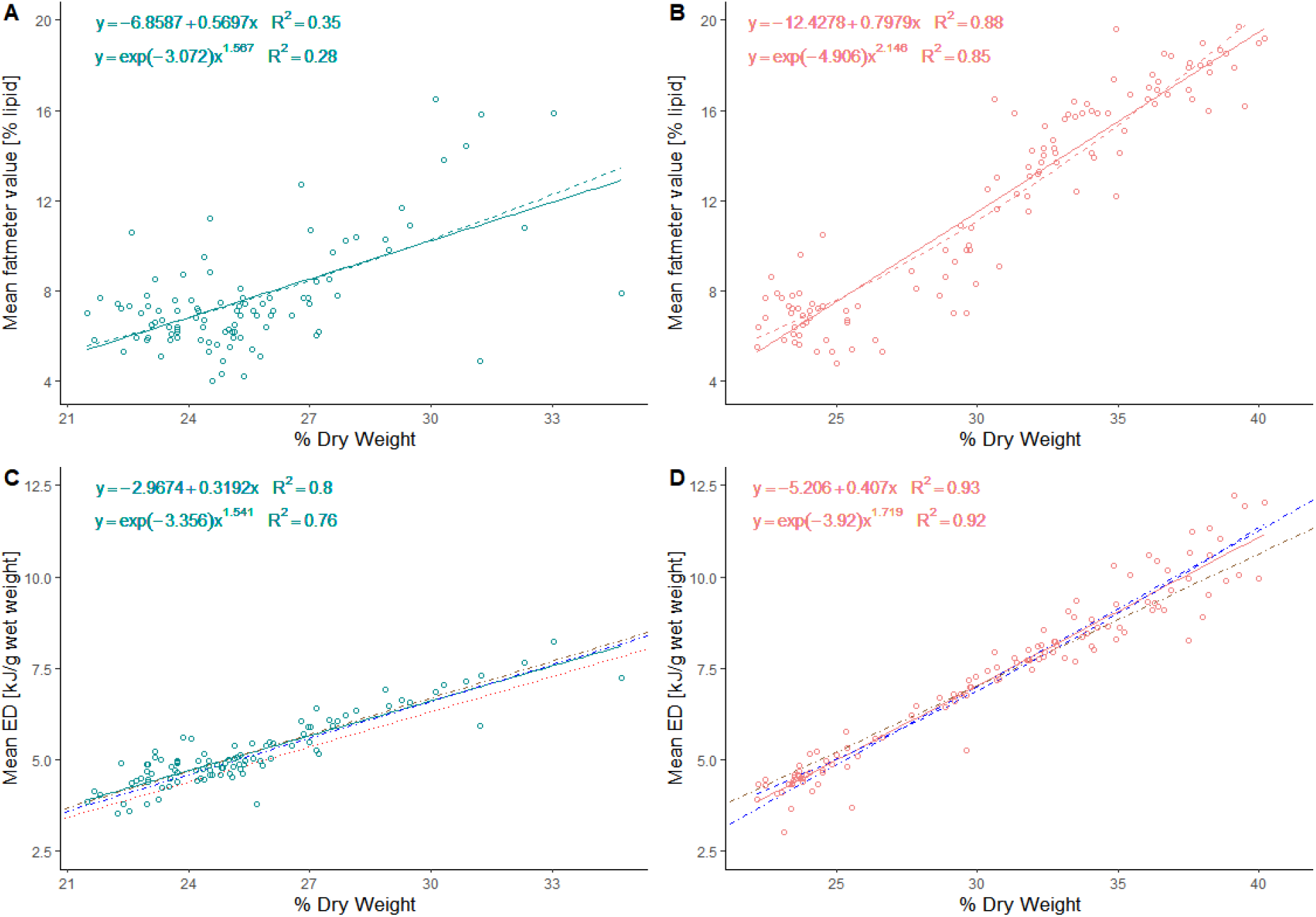
A) Fatmeter values (% lipid) against dry weight (g) of anchovy: linear model (solid line), log-log model (dashed line). B) Fatmeter values (% lipid) against dry weight (g) of sardine: linear model (solid line), log-log model (dashed line). C) Energy density in wet weight basis (kJ/g wet weight) against dry weight (g) of anchovy: linear model (cyan solid line) with equation, log-log model (cyan solid line) with equation, Gatti’s linear model (blue dot-dash line), Gatti’s log-log model (blue dashed line), Tirelli’s linear model (red dotted line) and Albo linear model (brown dot-dash line). D) Energy density in wet weight basis (kJ/g wet weight) against dry weight (g) of sardine: linear model (pink solid line) with equation, log-log model (pink dashed line) with equation, Gatti’s linear model (blue dot-dash line), Gatti’s log-log model (blue dashed line) and Albo linear model (brown dot-dash line).

With respect to fatmeter-DW relationship (Table 2), linear and log-log models displayed a strong fit for sardine (R^2^ of 0.88 and 0.85 for linear and log-log models, respectively), while a rather scarce fit was observed for anchovy (R^2^ values 0.35 and 0.28 for linear and log-log models, respectively) (Figure 5A-B).

**Table 2.** Coefficients of both linear and log-log regression models between fatmeter values (% lipid) and dry weight (%).

| Fatmeter vs DW | Species | R<sup>2</sup> | Coefficient | Estimate | Std. error |
| --- | --- | --- | --- | --- | --- |
| Linear model | Anchovy | 0.351 | Intercept | -6.859 | 1.994 |
|  |  |  | Slope | 0.570 | 0.078 |
|  | Sardine | 0.796 | Intercept | -10.571 | 1.060 |
|  |  |  | Slope | 0.741 | 0.034 |
| Log-log model | Anchovy | 0.282 | Intercept | -3.073 | 0.810 |
|  |  |  | Slope | 1.567 | 0.251 |
|  | Sardine | 0.849 | Intercept | -4.906 | 0.283 |
|  |  |  | Slope | 2.146 | 0.083 |

Conversely, the ED-DW models exhibited a very strong fit for both species being R^2^ values from 0.76 to 0.80 for anchovy and 0.92-0.93 for sardine (Figure 5C-D). Moreover, sardine models showed a slightly steeper slope, therefore the same rise in DW meant a higher increment in ED for sardine than for anchovy.

All the pair-wise Spearman correlations between condition indices were significantly positive (p<0.05) for anchovy. Such correlations were stronger in the resting period compared to the reproductive period (Figure 6 A-C).

**Figure 6.**
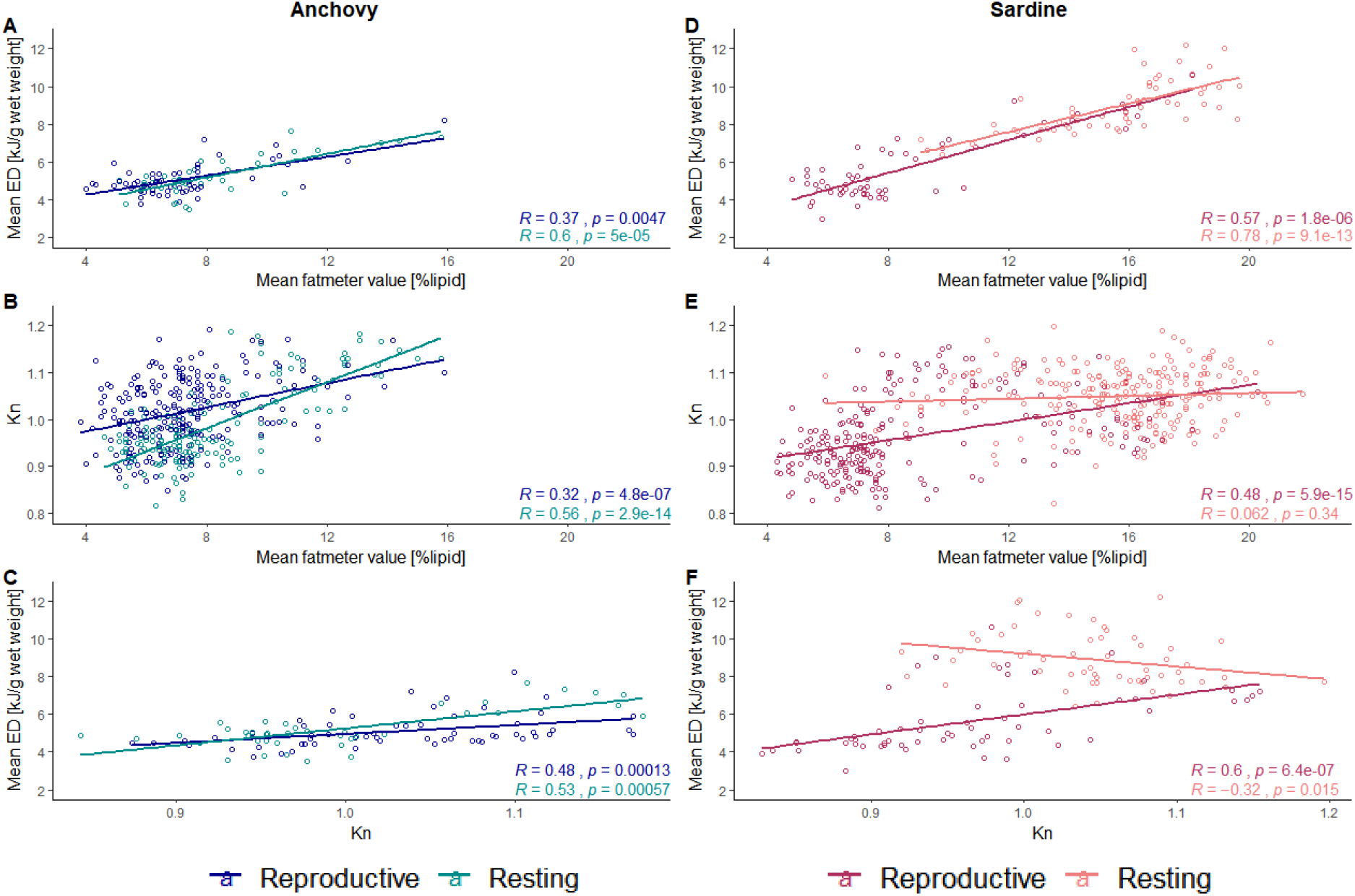
Spearman correlation tests between each of the condition indices for anchovy (A-C) and sardine (D-F) during the resting and reproductive period.

According to the Spearman correlation test, there was a significant relationship between every condition index in both resting and reproductive period for sardine (p-value < 0.05), except for Kn and fatmeter during the resting period (p = 0.34) (Figure 6 D-F). As opposed to the other pair-wise tests, Kn and ED of sardine during resting period exhibited a negative correlation (R = −0.32). With respect to the other comparisons, a stronger correlation was observed between fatmeter and ED in both periods.

Anchovy’s linear regression model between fatmeter values and ED exhibited a significantly better fit (R^2^ = 0.49) than the log-log model (R^2^ = 0.38) (Figure 7A). Linear model describing the relationship between mean fatmeter values and mean ED for sardine provided a slightly better fit (R^2^ = 0.82) than the log-log model (R^2^ = 0.81) (Figure 7B).

**Figure 7.**
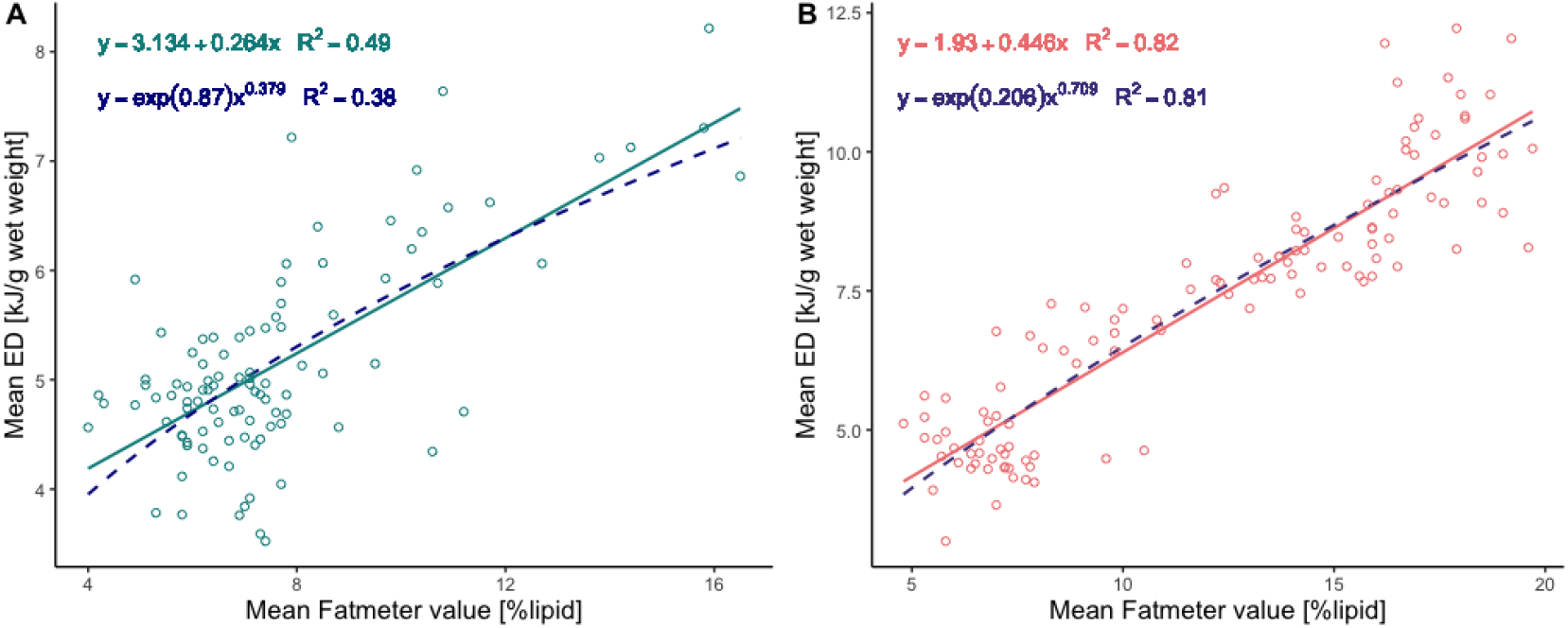
Energy density in wet weight basis (kJ/g ww) against mean fatmeter values (% lipid) of A) anchovy and B) sardine. Linear model (solid line), log-log model (dashed line).

## 4. Discussion

The key aim of this study was to assess the suitability of fish fatmeter as a surrogate of bomb calorimetry for sardine and anchovy. Since our results have revealed that fatmeter and bomb calorimetry measurements are highly correlated for both species, fatmeter can be used to estimate the energy content of sardine and anchovy in the Northwestern Mediterranean Sea.

### 4.1. Monthly variation in indirect and direct condition indices

Tracking monthly variation in energy content can help in predicting the response of small pelagic fish to environmental changes (Albo-Puigserver et al., 2020), thus choosing the condition index that better reflects the actual energy content is crucial.

Consistently with previous body condition assessments of anchovy and sardine from the southern Catalan coast (Albo-Puigserver et al., 2020, 2017), our results showed that year-round variation in body condition indices was more pronounced for sardine than for anchovy. This interspecies difference in body condition seasonal patterns reflects their distinct life-history strategies. Indeed sardine behaves as capital breeder (Albo-Puigserver et al., 2020, 2017; Brosset et al., 2015b; Ganias et al., 2007; Nunes et al., 2011), accumulating energy storage during the resting period when high fatmeter and ED values were recorded. While ED values of sardine had been increasing during summer, fatmeter values started to slowly decrease as lipids are generally the first macromolecule to be catabolized for energy in fishes (Shulman and Love, 1999; Tocher, 2003). Then the significant drop in all, indirect and direct, indices recorded from November to February represents the overall weight loss and lipid depletion in muscle, compensating for the energy invested for gonad development (Brosset et al., 2015b).

Interestingly, the high lipid % and ED values recorded for sardine in summer and at the beginning of autumn, with ED values close to 10 kJ·g^−1^ww, contrasted with the lower values previously reported in the southern Catalan coast in 2012 (maximum ED values close to 7 kJ·g^−1^ww; Albo-Puigserver et al., 2020). Low body condition and energy reserves of sardine, before the reproduction period, have been related to unfavorable environmental conditions and changes in primary and secondary production (Brosset et al 2015b; Albo-Puigserver et al., 2020). Therefore, the high energy content of sardine observed in the present study may result from better environmental conditions that could have contributed to a higher reproduction success (Garrido et al., 2007). However, further analysis combining condition indices and environmental variables should be performed to investigate this further.

On the other hand, the beginning of anchovy reproductive period is coupled with the late-winter peak in phytoplankton biomass and to the subsequent rise in zooplankton abundance (Palomera et al., 2007). As such, energy investment for reproduction could be partially counterbalanced by food intake thanks to the high prey availability (Basilone et al., 2006; Estrada, 1996). This explain the lower values and the less marked variation in energy content observed for anchovy in comparison with sardine and illustrates the income breeder behavior observed for this species.

### 4.2. Variation in energy content with length, sex, and season

In this study, the evolution of fatmeter values and ED with size displayed different seasonal patterns between sardine and anchovy. With respect to anchovy, high fatmeter values were recorded for some small individuals. This could be due to the fact that smaller individuals were measured in couple; thus, the related fatmeter values may be hindered and represent incorrect measurements of the lipid content. As far as sardine are concerned, the low fatmeter values recorded for large-class individuals in summer and winter may derive from a spawning period starting earlier and lasting longer, as previously observed by Nunes (2011). As there is not much literature on fatmeter values variation with length, we encourage future research to cover this aspect.

Despite previous studies analyzing ED variation with size reported that their relationship was not always linear (Pedersen and Hislop, 2001; Tirelli et al., 2006), here, anchovy and sardine ED-size models exhibited a positive linear fit with increasing length. These results contrast with the decrease in ED observed for larger anchovy individuals in Gatti et al. (2018) and Tirelli et al. (2006). However, it should be considered that in this study, only few individuals belonging to the larger size-classes (> 15 cm) were sampled, whereas in Gatti et al. (2018) larger individuals reached 18 and 25 cm for anchovy and sardine, respectively. This is also true for sardine as Gatti et al. (2018) reported a drop in ED for individuals over 19 cm, however our sample did not include fish longer than 18 cm. The paucity of larger-class individuals have already been attested for sardine populations from the NW Mediterranean (Albo-Puigserver et al., submitted; Pennino et al., 2020b, 2020a; Saraux et al., 2019; Van Beveren et al., 2014) and the Bay of Biscay (Véron et al., 2020).

While the season factor significantly affected both condition indices for both species, the effect of sex was limited and therefore not included in most models. Indeed, for anchovy, sex had a significant influence on fatmeter values, but not on ED. Conversely, sardine fatmeter values did not vary between males and females, in accordance with previous observation on muscle lipid content by Garrido et al. (2008). However, when relevant, the impact of sex for both species could be traced mainly to the presence of immature individuals (indeterminates); instead, differences in sex among adult individuals, i.e. males and females, was significant only in winter for both sardine and anchovy.

### 4.3. Validating fatmeter as surrogate of energy density

Even though the accuracy and repeatability of fatmeter analysis for sardine and anchovy has already been assessed (Brosset et al., 2015a), this study presents the validation of fatmeter analysis as a proxy of bomb calorimetry. The relationship between fatmeter and DW was investigated for both species using both linear and log-log models. Fatmeter-DW models can be used to obtain an estimation of lipid content not only in body condition assessments for ecological purposes, but also for fish meal production and human consumption (Albrecht-Ruiz and Salas-Maldonado, 2015; Marin et al., 2010; Šimat and Bogdanović, 2012; Tufan et al., 2011; Zlatanos and Laskaridis, 2007). Anchovy displayed a stronger relationship between ED and DW than between fatmeter and DW, whereas for sardine both regressions results were strong. In any case, ED and DW were highly correlated for both species (R^2^ > 0.75). Although Gatti et al. (2018) suggested that ED-DW models are species- and area-specific, ED-DW models from this study were similar to the ones fitted to populations from different geographical areas (Table S4), i.e. the Bay of Biscay and the English Channel (Gatti et al., 2018), Tarragona (Albo-Puigserver et al., 2020; unpub. data) and the Adriatic Sea (Tirelli et al., 2006). Nevertheless, Welch’s test results (Table S5) revealed that the coefficient estimates from our study were significantly different to the ones from previous literature except for the slope of anchovy’s linear model from Tirelli et al. (2006). However, the statistical difference may not reflect an actual ecological difference as overall the regression lines were very similar. A previous study suggested the use of ED-DW log-log models, instead of linear models, as they may better account for ontogenetic variability (Gatti et al., 2018a). Despite this, in our study, linear models provided a slightly better fit than log-log models for both species and this may be explained by the clear prevalence of adults, who were presumably mature, and the absence of early juveniles in our samples.

According to Spearman’s correlation test, positive strong correlations were observed between fatmeter and ED. Furthermore, the results highlighted differences in all the pairwise-correlations between condition indices depending on the reproductive state of the fish as previously observed by Brosset et al. (2015). Indeed, fatmeter and ED for anchovy and sardine clearly showed higher correlation during the resting period, when the lipid content is higher, compared to the reproductive period. Neither sardine’s ED nor fatmeter values were significantly correlated with Kn during the resting period. Therefore, future body condition assessments for this species should consider that Kn may not be a proper condition index during non-reproductive period, as it may not reflect the energetic status.

ED-fatmeter linear regression models provided a weaker fit for anchovy than for sardine. The stronger ED-fatmeter relationship for sardine may be traceable to its higher energy reserves, consisting mainly of lipids, which are the main driver of ED variability (Albo-Puigserver et al., 2020; Rosa et al., 2010). In contrast, anchovy variation in ED is probably prompted by changes in other macromolecules, i.e. proteins, which cannot be measured with fatmeter analysis (Albo-Puigserver et al., 2020). Indeed, while direct calorimetry estimates the energy density of the whole proximate composition (i.e. lipids, proteins, and carbohydrates), fatmeter gives an indirect measure only of the lipid content.

Previous study by Brosset et al. (2015) validated the use of fatmeter comparing its lipid estimations with the ones by thin-layer chromatography with Iatroscan MK-VI. In contrast with our results, the relationship between lipid content estimations by means of fatmeter and chromatography was stronger for anchovy than for sardine. This weaker relationship for sardine may be attributable to the fact that chromatography analysis were performed only on gonad and muscle tissue, without considering mesenteric fat, while fatmeter measurements were taken on the whole fish. Indeed mesenteric fat is a long-term storage site for sardine (Lloret et al., 2014) and it tends to increase from spring to summer in concomitance with the rise in zooplankton abundance in the study area (Albo-Puigserver et al., 2017). Furthermore, a previous study on Atlantic herring, *Clupea harengus*, evidenced a strong correlation between fatmeter measurements and mesenteric fat (McPherson et al., 2011).

Overall, the relative ability of fatmeter values to estimate energy content proved to be significantly higher for sardine. This outcome can be traced to the smaller size of anchovy that increases the difficulty and reduces the repeatability of fatmeter measurements (Brosset et al., 2015a).

The interchangeability of the two techniques, fatmeter and calorimetry analysis, may lead to an implementation of fatmeter measurements in the body condition assessment protocol. Indeed, ED-fatmeter models could provide adequate estimates of energy density in field applications of bioenergetics models.

## 5. Conclusions

Our study supports the use of fatmeter analysis as a surrogate of bomb calorimetry to infer the energy density of both sardine and anchovy. Thanks to the fatmeter validation provided in this study, future bioenergetic assessments of these two species could benefit from this technique as it allows rapid energy content estimation and can be carried out *in situ*. In turn, this would shorten the time required for large-scale and continuous monitoring that contribute to the understanding of population dynamics and to fish stock management and conservation (Rosa et al., 2010). Moreover, as opposed to Kn values, which are population-specific (Lloret et al., 2014), fatmeter values may be compared among populations from different geographical areas.

However, our results confirm that the validation of body condition indices should consider the ontogenetic variation (Schloesser and Fabrizio, 2017; Wuenschel et al., 2006), the reproductive state (Brosset et al., 2015b; Nunes et al., 2011) and the seasonality. Indeed, fatmeter correlation with ED was demonstrated to be stronger for sardine than for anchovy and it changed depending on the reproductive period. Similarly, fatmeter values and dry weight displayed stronger relationship for sardine. Therefore, fatmeter analysis could conceivably be a better proxy of energy content for species that store high quantity of fat such as sardine; nevertheless, its use proved to be appropriate also for anchovy, in particular during the resting period and for larger individuals.

Finally, using the species-specific equations provided in this study, fatmeter values could be transformed into ED units and eventually serve as inputs for bioenergetics models.

## AUTHOR CONTRIBUTIONS

Claudia Campanini: Conceptualization, Investigation, Formal analysis, Writing – Original Draft, Visualization

Marta Albo-Puigserver: Conceptualization, Investigation, Writing – Review & Editing, Supervision Sara Gérez: Investigation

Elena Lloret-Lloret: Investigation, Data curation, Writing – Review & Editing

Joan Giménez: Investigation, Writing – Review & Editing

Maria Grazia Pennino: Formal analysis, Writing – Review & Editing

Jose Maria Bellido: Writing – Review & Editing

Ana I. Colmenero: Investigation, Writing – Review & Editing

Marta Coll: Conceptualization, Writing – Review & Editing, Supervision

## Acknowledgments

This work was supported by PELWEB [ES-PN-2017-CTM2017-88939-R] project. C.C. was funded by an Erasmus+ Mobility for Traineeships scholarship. We would like to thank the Cofraria de Pescadores de Barcelona for their help in the acquisition of the samples.

