## Supplementary materials for "Energy content of anchovy and sardine using surrogate calorimetry methods"

***Variation in energy content with length, sex, and season***

***Table S1.*** Numerical summary of the best Generalized Additive Model (GAM) obtained for A) Fatmeter for anchovy, B) ED for anchovy, C) Fatmeter for sardine, D) ED for sardine. Chosen models in bold. Covariates’ acronym: ID = individual random effect. Statistic acronyms: D %= explained deviance in percentage; AIC = Akaike information criterion. Note that s(covariable) means that such variable has been introduced in the model through a spline function and k stands for the number of knots set for the spline function.

| **A) FATMETER GAM FOR ANCHOVY**  Family: gaussian  Link function: identity | | | |
| --- | --- | --- | --- |
| **N.** | **Covariates** | **D %** | **AIC** |
| 1 | s(Total_Length, k=6) + s(Total_Length, by=Season,k=6) + s(ID, bs="re") + Season + Sex + Sex*Season | 36.7% | 1608.027 |
| **2** | **s(Total_Length, k=6) + s(Total Length, by=Season,k=6) + Season +s(ID, bs="re") + Sex** | **34.3%** | **1609.962** |
| 3 | s(Total_Length, k=6) + s(Total Length, by=Season, k=6) +s(ID, bs="re") + Season | 32.6% | 1616.738 |
| 4 | s(Total_Length, k=6) + s(ID, bs="re") + Season + Sex | 27.4% | 1639.539 |
| 5 | s(Total_Length, k=6) + s(Total Length, by=Season, k=6) + Season + Sex | 27.2% | 1647.815 |
| 6 | s(Total_Length, k=6) + s(Total Length , by=Season, k=6) + Season | 26.7% | 1647.568 |
| 7 | s(Total_Length, k=6) + Season + Sex | 22.6% | 1662.162 |
| 8 | s(Total_Length, k=6) + Season | 22% | 1661.100 |
| 9 | s(Total_Length, k=6) + s(Total Length, by=Season, k=6) | 16.8% | 1690.034 |
| 10 | s(Total_Length, k=6) | 10.3% | 1710.705 |
| **B) ED GAM FOR ANCHOVY**  Family: gaussian  Link function: identity | | | |
| **N.** | **Covariates** | **D %** | **AIC** |
| 1 | s(Total_Length, k=6) + s(Total_Length, by=Season, k=6) + s(ID, bs="re") + Season + Sex + Sex*Season | 55% | 216.1267 |
| 2 | s(Total_Length, k=6) + s(Total_Length, by=Season, k=6) + s(ID, bs="re") + Sex + Season | 52.9% | 212.7637 |
| **3** | **s(Total_Length, k=6) + s(Total_Length, by=Season, k=6) + s(ID, bs="re") + Season** | **52.7%** | **209.1886** |
| 4 | s(Total_Length, k=6) + s(Total_Length, by=Season, k=6) + Season | 46.5% | 219.4070 |
| 5 | s(Total_Length, k=6) + s(ID, bs="re") + Season | 37.8% | 233.0843 |
| **C) FATMETER GAM FOR SARDINE**  Family: gaussian  Link function: identity | | | |
| **N.** | **Covariates** | **D %** | **AIC** |
| 1 | s(Total_Length, k=6) + s(Total_Length, by=Season, k=6) + s(ID, bs="re") + Season + Sex + Sex*Season | 75.4% | 2218.111 |
| **2** | **s(Total_Length, by=Season, k=6) + s(ID, bs="re") + Season** | **74.1%** | **2229.661** |
| 3 | s(Total_Length, by=Season, k=6) + Season | 72.5% | 2255.418 |
| 4 | s(Total_Length, k=6) + s(ID, bs="re") + Season + Sex*Season | 71.7% | 2272.773 |
| 5 | s(Total_Length, k=6) + s(ID, bs="re") + Season + Sex | 71.4% | 2271.437 |
| 6 | s(Total_Length, k=6) + s(ID, bs="re") + Season | 71.2% | 2272.029 |
| 7 | s(ID, bs="re") + Sex*Season + Sex + Season | 65.8% | 2354.725 |
| **D) ED GAM FOR SARDINE**  Family: gaussian  Link function: identity | | | |
| **N.** | **Covariates** | **D %** | **AIC** |
| 1 | s(Total_Length, k=6) + s(Total_Length, by=Season, k=6) + s(ID, bs="re") + Sex + Season + Sex*Season | 74% | 393.7332 |
| 2 | s(Total_Length, by=Season, k=6) + Season + Sex | 72.1% | 396.1380 |
| 3 | s(Total_Length, by=Season, k=6) + s(ID, bs="re") + Season | 72.1% | 397.5740 |
| **4** | **s(Total_Length, k=6) + Season + Sex** | **69,2%** | **398.0542** |
| 5 | Season + Season*Sex | 67.9% | 406.0001 |

***Table S2.*** Summary of Fatmeter-size (a) and ED-size (b) GAMs for anchovy. Asterisks (*) next to the p-values indicate statistical significance.

| **a) FATMETER**  Fatmeter ~ s(Total_Length, k=6) +s(Total_Length, by=Season, k=6) + Season + s(ID, bs="re") + Sex | | **Anchovy** | | |
| --- | --- | --- | --- | --- |
|  |  | Estimate | Std. error | P-value |
| Fixed effect: Season | Summer  Autumn  Winter  Spring | 7.672  1.773  3.783  5.330 | 0.247  0.418  0.445  0.653 | < 2e-16*  2.75e-05*  4.20e-16*  5.17e-15* |
| Fixed effect: Sex | I  M | -1.481  -0.490 | 0.659  0.202 | 0.025*  0.016* |
| Smoothed length effect | Total length | 1.38e-08* | | |
| Smoothed length effect by season | Summer  Autumn  Winter  Spring | <0.001*  0.017*  0.077  0.997 | | |
| Smoothed random individual effect | ID | 1.16e-09* | | |
| R^2^_adj_ | | 0.32 | | |
| AIC | | 1610 | | |
| **b) ED**  ED ~ s(Total_Length, k = 6) + s(Total_Length, by = Season, k = 6) + Season + s(ID, bs = "re") | | **Anchovy** | | |
|  |  | Estimate | Std. error | P-value |
| Fixed effect: Season | Summer  Autumn  Winter  Spring | 4.909  0.695  1.359  2.498 | 0.154  0.276  0.311  0.452 | < 2e-16*  0.014*  3.24e-05*  3.12e-07* |
| Smoothed length effect | Total length | 1.56e-09* | | |
| Smoothed length effect by season | Summer  Autumn  Winter  Spring | 2.14e-06*  2.23e-05*  ≈ 1  0.067* | | |
| Smoothed random individual effect | ID | 0.001* | | |
| R^2^_adj_ | | 0.49 | | |
| AIC | | 219 | | |

***Table S3.*** Summary of Fatmeter-size (a) and ED-size (b) GAMs for sardine. Asterisks (*) next to the p-values indicate statistical significance.

| **a) FATMETER**  Fatmeter.mean ~ Season + s(ID, bs="re") + s(Total_Length, k = 6) | | **Sardine** | | |
| --- | --- | --- | --- | --- |
|  |  | Estimate | Std. error | P-value |
| Fixed effect: Season | Summer  Autumn  Winter  Spring | 17.190  -5.136  -4.322  4.479 | 0.333  0.529  0.857  1.205 | < 2e-16*  < 2e-16*  6.6e-07* <0.001* |
| Smoothed length effect by season | Summer  Autumn  Winter  Spring | 2.45e-07*  < 2e-16*  0.388  4.52e-15* | | |
| Smoothed random individual effect | ID | 1.38e-07* | | |
| R^2^_adj_ | | 0.74 | | |
| AIC | | 2229 | | |
| **b) ED**  ED ~ s(Total_Length, k = 6) + Season + Sex | | **Sardine** | | |
|  |  | Estimate | Std. error | P-value |
| Fixed effect: Season | Summer  Autumn  Winter  Spring | 9.786  -3.481  -3.995  -1.487 | 0.293  0.350  0.370  0.354 | < 2e-16*  < 2e-16*  < 2e-16*  5.41e-05* |
| Fixed effect: Sex | M-F | -0.514 | 0.253 | 0.045* |
| Smoothed length effect | Total length | 0.0051* | | |
| R^2^_adj_ | | 0.68 | | |
| AIC | | 398 | | |

***Fatmeter validation***

***Table S4.*** Coefficients of both linear and log-log regression models between energy density (ED) expressed in wet weight basis (kJ/g of wet weight) and dry weight (DW) from this study, from published studies: Gatti et al., 2018, Tirelli et al., 2006 and from Albo-Puigserver et al. 2020; unpub. data.

| ED against DW | Species | R^2^ | Coefficient | Estimate | Std. error |
| --- | --- | --- | --- | --- | --- |
| Linear models | Anchovy | 0.8004 | Intercept  Slope | -2.967  0.319 | 0.410  0.016 |
|  | Sardine | 0.9201 | Intercept  Slope | -5.206  0.407 | 0.303  0.010 |
| Log-log models | Anchovy | 0.7672 | Intercept  Slope | -3.356  1.541 | 0.277  0.086 |
|  | Sardine | 0.9201 | Intercept  Slope | -3.920  1.719 | 0.158  0.046 |
| Linear models  from Gatti et al., 2018 | Anchovy | 0.894 | Intercept  Slope | - 3.403  0.333 | 0.254  0.009 |
|  | Sardine | 0.940 | Intercept  Slope | −5.85  0.428 | 0.226  0.008 |
| Log-log models  from Gatti et al., 2018 | Anchovy | 0.907 | Intercept  Slope | −3.302  1.523 | 0.127  0.038 |
|  | Sardine | 0.942 | Intercept  Slope | −3.995  1.740 | 0.101  0.030 |
| Linear model  From Tirelli et al., 2006 | Anchovy | 0.82 | Intercept  Slope | -3.31691 0.32101 | /  / |
| Linear models  from Albo-Puigserver  (unpub. Data) | Anchovy | 0.91 | Intercept  Slope | -3.3908  0.3362 | 0.308  0.012 |
|  | Sardine | 0.94 | Intercept  Slope | -3.8155  0.3611 | 0.266  0.010 |

***Table S5.*** Results of Welch’s two sample t-test between coefficients of both linear and log-log regression models between energy density expressed in wet weight basis (kJ/g of wet weight) and dry weight from this study, from Gatti et al., 2018, Tirelli et al., 2006 and Albo-Puigserver at al., 2020; unpub. data. Asterisks (*) next to the t-values and the degrees of freedom (df) indicate statistical significance.

| **Our models** |  | **Gatti et al., 2018** | **Albo-Puigserver et al 2020,**  **unpub. data** | **Tirelli et al., 2006** |
| --- | --- | --- | --- | --- |
| Anchovy linear model | Intercept | t = 90.315  df = 16766 | t = 82.299*  df = 18538* | t = 32.359*  df = 13328* |
|  | Slope | t = -76.523  df = 15702 | t = -85.773*  df = 18452* | t = -1.734  df = 10004 |
| Anchovy  log-log model | Intercept | t = -18.702  df = 14030 | /  / | /  / |
|  | Slope | t = 17.845  df = 13747 | /  / | /  / |
| Sardine  linear model | Intercept | t = 171.35  df = 1846 | t = -339.74*  df = 19665* | /  / |
|  | Slope | t = -164.69  df = 19177 | t = 325.17*  df = 19997* | /  / |
| Sardine  log-log model | Intercept | t = 40.246  df = 17009 | /  / | /  / |
|  | Slope | t = -36.457  df = 17170 | /  / | /  / |

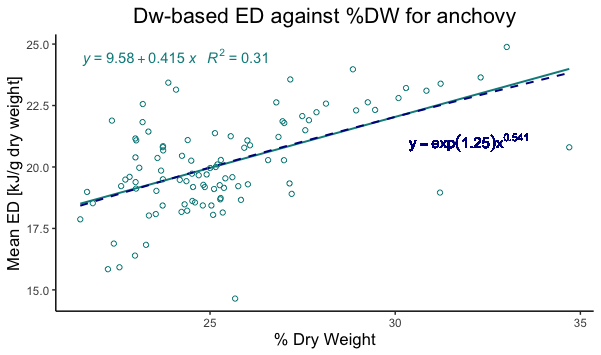

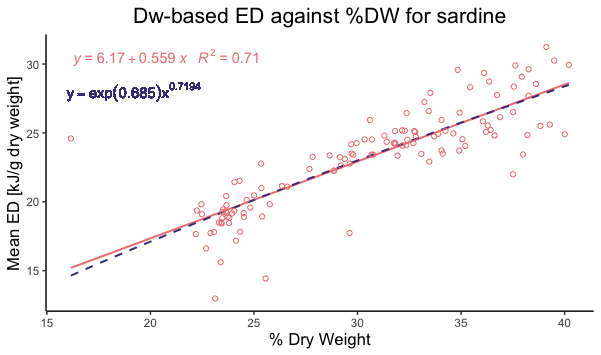

***Figure S1.*** Energy density in dry weight basis (Mean ED, kJ / g dry weight) against % dry weight (g) of anchovy (left) and sardine (right). Plotted data with regression models lines and equations: linear model (solid line) and log-log model (dashed line).
